# The worldwide invasion of *Drosophila suzukii* is accompanied by a large increase of transposable element load and a small number of putatively adaptive insertions

**DOI:** 10.1101/2020.11.06.370932

**Authors:** Vincent Mérel, Patricia Gibert, Inessa Buch, Valentina Rodriguez Rada, Arnaud Estoup, Mathieu Gautier, Marie Fablet, Matthieu Boulesteix, Cristina Vieira

## Abstract

Transposable Elements (TEs) are ubiquitous and mobile repeated sequences. They are major determinants of host fitness. Here, we portrayed the TE content of the spotted wing fly *Drosophila suzukii*. Using a recently improved genome assembly, we reconstructed TE sequences *de novo*, and found that TEs occupy 47% of the genome and are mostly located in gene poor regions. The majority of TE insertions segregate at low frequencies, indicating a recent and probably ongoing TE activity. To explore TE dynamics in the context of biological invasions, we studied variation of TE abundance in genomic data from 16 invasive and six native populations (of *D. suzukii*). We found a large increase of the TE load in invasive populations correlated with a reduced Watterson estimate of genetic diversity 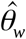 a proxy of effective population size. We did not find any correlation between TE contents and bio-climatic variables, indicating a minor effect of environmentally induced TE activity. A genome-wide association study revealed that *ca*. 5,000 genomic regions are associated with TE abundance. We did not find, however, any evidence in such regions of an enrichment for genes known to interact with TE activity (e.g. transcription factor encoding genes or genes of the piRNA pathway). Finally, the study of TE insertion frequencies revealed 15 putatively adaptive TE insertions, six of them being likely associated with the recent invasion history of the species.

## Introduction

Transposable Elements (TEs) are selfish genetic elements. Despite being mostly neutral or deleterious, they persist and proliferate in populations by copying and pasting themselves in genomes (Doolittle and Sapienza 1980; Orgel and Crick 1980; Charlesworth and Charlesworth 1983). The interest for those sequences considerably rose in the 2000’s, with the discovery of some TE insertions having a functional, and potentially adaptive, effect on their host (Mi et al. 2000; Daborn et al. 2002; Niu et al. 2019). The parallel completion of the first sequencing projects confirmed TE ubiquity and largely contributed to the growing interest for such sequences (C. elegans Sequencing Consortium 1998; 2000; Lander et al. 2001; 2002; Schnable et al. 2009).

The nature and intensity of TE deleterious effects may vary with their genomic localization (Mérel et al. 2020). First, TEs close to genes can alter their function. Second, TEs in highly recombining regions, are more likely to promote ectopic recombination, *i*.*e*. recombination between more-or-less identical sequences inserted at different locations in the genome. Third, recessive deleterious TEs are more likely to impact fitness when located on a chromosome in a hemizygous state (*e*.*g*. the X chromosome in males in a XY sex determination system). The strength of selection acting against TEs hence depends on the genomic region and may result in a local variation of TE density. In agreement with such expectations, TE density was found to be negatively correlated with gene density and local recombination rate in several species (Boissinot et al. 2001; Bartolomé et al. 2002). On the other hand, studies focusing on the *D. melanogaster* genome did not reveal a systematic lower TE content on the X-chromosome, which is hemizygous in males (Kofler et al. 2012; Cridland et al. 2013).

TE insertion frequencies reflect both TE activity and the selection acting upon them. Low frequency TE insertions are likely to be recent, or strongly selected against, or both. Conversely, high frequency TE insertions are likely to be old and only weakly subjected to purifying selection. As mentioned previously, TEs that are in the vicinity of genes and/or located in highly recombining regions are expected to be selected against. Accordingly, TE insertion frequencies were found to be negatively correlated with recombination rate and distance to the nearest gene in *D. melanogaster* (Kofler et al. 2012). In Drosophila, the overall distribution of TE frequencies seems compatible with an active repeatome (Kofler, Nolte, et al. 2015; Hill 2019) For example 80% of the insertions have a frequency lower than 0.2 in *D. melanogaster* and its close relative *D. simulans (Kofler, Nolte, et al. 2015)*.

Between population variation of TE content has been reported in various intraspecific studies. So far, the factors underlying such differences remain unclear. The effective population size (N_e_) may play a prominent role in modulating TE contents. Considering that TEs are mostly deleterious, and that small N_e_ leads to a less efficient purifying selection, small N_e_ should be associated with high TE content (Lynch and Conery 2003). In support for this hypothesis Lynch & Connery (2003) found a significant correlation between genome size and estimates of the scaled mutation rate θ=N_e_ *μ* (with µ the mutation rate) across populations representative of various species. At the intraspecific level, if a higher TE content in some populations has sometimes been suggested to result from a reduction of their N_e_ (García Guerreiro et al. 2008; García Guerreiro and Fontdevila 2011; Talla et al. 2017), to our knowledge the above expected correlation has not been reproduced at this evolutionary scale. Variation in TE content may also rely on changes in TE activity in relation with the environment (Vieira et al. 1999; Stapley et al. 2015). In Drosophila, several laboratory experiments suggest that TE activity may respond to the environment (García Guerreiro 2012; Horváth et al. 2017), but *in natura* studies considering the whole repeatome remain rare and a possible confounding effect of the demographic history cannot be excluded (Lerat et al. 2019). Finally, the host genotype may explain intraspecific variation of TE abundance. For instance, in Drosophila, several studies found different levels of activity among isogenic lines (Biémont et al. 1987; Pasyukova and Nuzhdin 1993; Díaz-González et al. 2011).

The study of intraspecific variations in TE content and the underlying determining factors is valuable as TEs may also be important for adaptation (Daborn et al. 2002; Van’t Hof et al. 2016; Niu et al. 2019). Although some TE insertions exhibit a strong signal of positive selection and have been thoroughly validated experimentally, only few studies aimed at identifying putatively adaptive insertions at a genome-wide level (González et al. 2008; Li et al. 2018; Rishishwar et al. 2018; Rech et al. 2019). In addition, most of these studies deal with *D. melanogaster* (González et al. 2008; González et al. 2010; Blumenstiel et al. 2014; Rech et al. 2019).The most comprehensive of these studies analyzed genomic data on 60 worldwide natural *D. melanogaster* populations and reported 57 to 300 putatively adaptive insertions (depending on the degree of evidence considered) among the ∼800 polymorphic insertions identified in the reference genome (Rech et al. 2019). Considering that approximately twice as many non reference TE insertions as reference insertions may segregate in a single population (Kofler et al. 2012), quite a high number of TE-induced adaptations is therefore expected. However, it remains unclear how important TEs are as substrates of adaptation considering the paucity of studies and their focus on reference genome insertions.

Invasive species provide a unique opportunity to study the combined effect of *in natura* N_e_ variations and environmental variations both on TE abundance and TE adaptive potential. Invasive populations often go through demographic bottlenecks allowing to test for an effect of N_e_ on TE abundance (Estoup et al. 2016). Individuals from invasive populations also encounter new environmental conditions, allowing to test for an effect of bio-climatic variables on TE abundance. Because of the need of colonizing individuals to adapt to new environmental conditions, biological invasions are often used to study rapid contemporary adaptation (Lavergne and Molofsky 2007; Rollins et al. 2015). Yet, the particular role of TEs in the rapid adaptation of invasive species remains speculative. In particular, TEs have been proposed to explain, at least in part, the paradox of invasive species, *i*.*e*. the successful adaptation to a new environment despite a reduced genetic diversity caused by small founder population sizes (Stapley et al. 2015; Estoup et al. 2016; Marin et al. 2020). In response to environmental changes, TE sequences may be recruited and affect the expression of nearby genes. Furthermore, if a higher activity of TE is induced in response to environmental changes, the insertions could thus result in genetic variation, and potentially beneficial alleles.

In this paper, we focused on the spotted wing fly *D. suzukii*, a close relative of *D. melanogaster*, displaying the highest reported TE content among Drosophila (Sessegolo et al. 2016). *D. suzukii* is native from Asia and has invaded independently the American and European continents where it was introduced probably in the late 2000’s (Fraimout et al. 2017). Using the recently released high-quality genome assembly Dsuz-WT3_v2.0 based on Long PacBio Reads (Paris et al. 2020), we constructed a *de novo* TE database and found that TE represented 47 % of the genome. We further assessed TE insertion frequencies and TE abundance in 22 worldwide populations representative of the native area (n=6) and of the two main invaded areas in Europe (n=8) and America (n=8). The study of TE frequencies showed that the repeatome is highly active in *D. suzukii*: 75% of insertion segregated at a frequency < 0.25. We found that the TE content was significantly higher in invasive populations and was correlated with a reduction of N_e_. Finally, controlling for population structure, a genome scan conducted on polymorphic TE insertions identified 15 putatively adaptive TE insertions.

## Results

### A highly repeated reference genome

We found that the high-quality *D. suzukii* assembly Dsuz-WT3_v2.0 of Paris et al. (2020) is characterized by a high TE content. Overall, 47.07 % of the reference assembly is annotated as repeated sequences (fig. 1*A*). In terms of genomic occupancy, LTR is the predominant TE order with more than 20% of the sequence assembly corresponding to these elements, then LINEs (8.77%), DNA elements (6.99%), and RC (6.95%). 4.07% of the assembly is occupied by unknown repeated sequences. At a lower hierarchical level, the three most represented superfamilies are *Gypsy, Helitron* and *Pao*, corresponding to 13.65%, 6.95% and 6.44% of the assembly, respectively (supplementary table S1). The average percentage of genomic occupancy per superfamily is 1.88%. Regarding TE copy numbers, the top three superfamilies are *Helitron, Gypsy* and *Pao* (56,493, 39,189 and 15,555 copies, respectively) (fig. 1*A*). The average number of copies per superfamily is 4,963.

**Figure 1:**
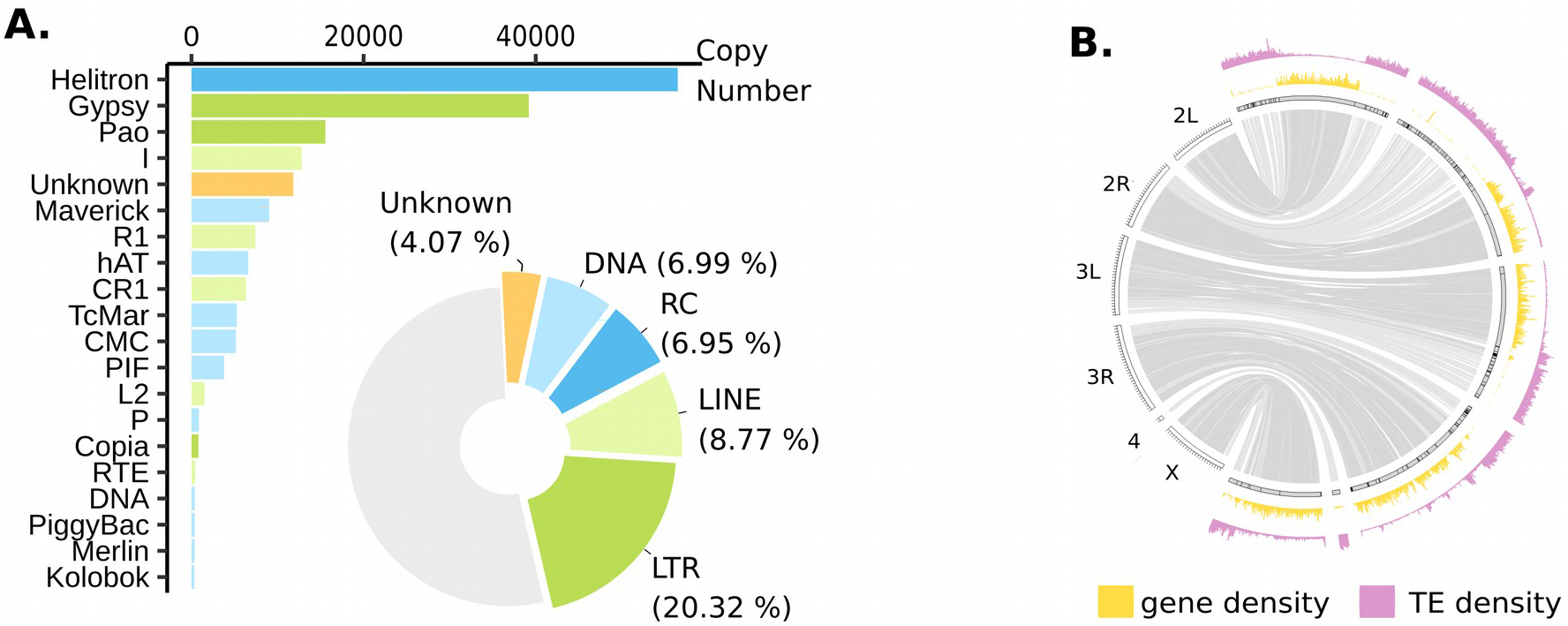
Main features of the TE content in the *D. suzukii* reference genome. A. TE copy numbers and TE genomic occupancy. Barplot representing TE copy numbers for the 20 TE superfamilies displaying the highest copy numbers Piechart illustrating genomic sequence occupancy of each TE order (in percentages of the assembly). Class I TEs are shown in green (light green for LINEs and darker green for LTR Elements). Class II TEs are shown in blue (light blue for DNA and darker blue for Rolling Circles (RC)). Non repeated sequences are shown in gray. B. Distribution of TEs and genes. TE density (pink outer graph) and gene density (yellow inner graph) are shown for windows of 200 kb. The maximum value of gene density is The maximum number of TE fragments is 713. Syntenic relationships with *D. melanogaster* assembly are shown inside using light links for regions of low gene density (< 7 genes per 200 kb) and dark links for regions of high gene density (>= 7 genes per 200 kb). Contigs are surrounded by black strokes. Ticks on *D. melanogaster* assembly are separated by one Mb.

Syntenic relationships with *D. melanogaster* genome have been established for 212 of the 546 contigs of *D. suzukii* assembly. A total of 241 Mb of the 268 Mb assembly have a clearly identified counterpart in the *D. melanogaster* genome (fig. 1*B*, supplementary table S2). Considering the observed bimodal distribution of gene density, we partitioned the *D. suzukii* assembly into gene-rich regions (≥ 7 genes per 200 kb; 121.8 Mb) and gene-poor regions (< 7 genes per 200 kb; 108 Mb) (fig 1*B*, supplementary fig. S1). TE fragment density also follows a bimodal distribution: 127.4 Mb correspond to TE-rich regions (≥ 165 TE fragments per 200 kb) and 102.4 Mb to TE-poor regions (< 165 TE fragments per 200 kb) (fig. 1*B*, supplementary fig. S2). TE-rich regions are enriched in gene-poor regions, and TE-poor regions are enriched in gene-rich regions (χ^2^ = 786.47, df = 1, p-value < 2.2×10^−16^). We did not find any difference in mean TE density between autosomal and X-linked contigs 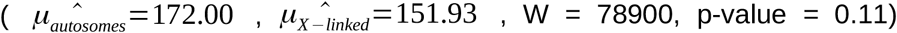. This conclusion holds when comparing autosomal and X-linked contigs as defined in Paris et al. (2020) using a female-to-male read mapping coverage ratio 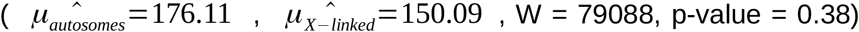. However, when considering only gene-rich regions, the mean TE density was far higher for X-linked contigs 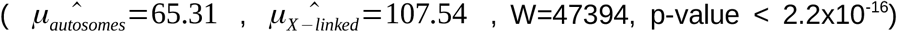. Once again, this conclusion holds when using autosomal and X-linked contigs as defined by Paris et al. (2020) 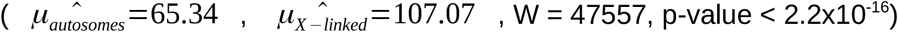.

### An active repeatome in the Watsonville reference population

The female used to establish the WT3 isofemale strain corresponding to the genome assembly was collected in Watsonville (CA, USA) (Paris et al. 2020). To thoroughly evaluate TE activity in this reference population, we assessed TE insertion frequencies in a PoolSeq sample of 50 *D. suzukii* individuals from Watsonville. Because TEs are mostly deleterious, rare TE insertions are likely to be recent insertions, not yet eliminated by selection, whereas fixed TE insertions are presumably old insertions weakly submitted to selection. It is worth stressing that, for the study of TE frequencies and abundances, we first used simulated PoolSeq data to validate our pipelines and to evaluate their performance and their sensibility to parameters such as sequencing coverage or number of individuals (see supplementary methods for details).

A total of 9,256 insertions were recovered in the reference population. The frequency distribution is approximately U-shaped (fig. 2*A*) with a majority of insertions segregating at low frequency (N = 6934, f < 0.25). 1,642 insertions are found at high frequency, in the reference population (f ≥ 0.75). Only a minority of insertions are of intermediate frequency (N = 680, 0.25 ≤ f < 0.75). Among the 654 families/pseudofamilies found in the whole dataset, 473 were present in the reference population. 102 belonged to the DNA order, 98 to the LINE order, 175 to the LTR order, 46 to the RC and 52 were Unknown. Only 119 TE families/pseudofamilies presented more than 10 insertions: 25 DNA families/pseudofamilies, 32 LINEs, 32 LTR, 6 RC and 24 Unknown. The vast majority of these families presented a median frequency lower than 0.25 (N = 80) (fig. 2*B*). Only four families displayed a median frequency between 0.25 and 0.75. Finally, 35 families had a median frequency superior or equal to 0.75. We did not find evidence that the number of TE families in these categories differed between TE orders (supplementary table S3; χ^2^ = 4.94, df = 8, p-value = 0.76). However, the mean frequency was slightly different (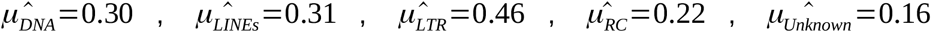, Kruskall-Wallis χ^2^ = 92.35, df = 4, p-value < 2.2×10^−16^). TE insertion frequencies were not evenly distributed along the assembly: mean TE insertion frequency was considerably lower in gene-rich windows. (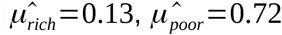, W = 18863, p-value < 2.2e-16; supplementary fig. S3).

**Figure 2:**
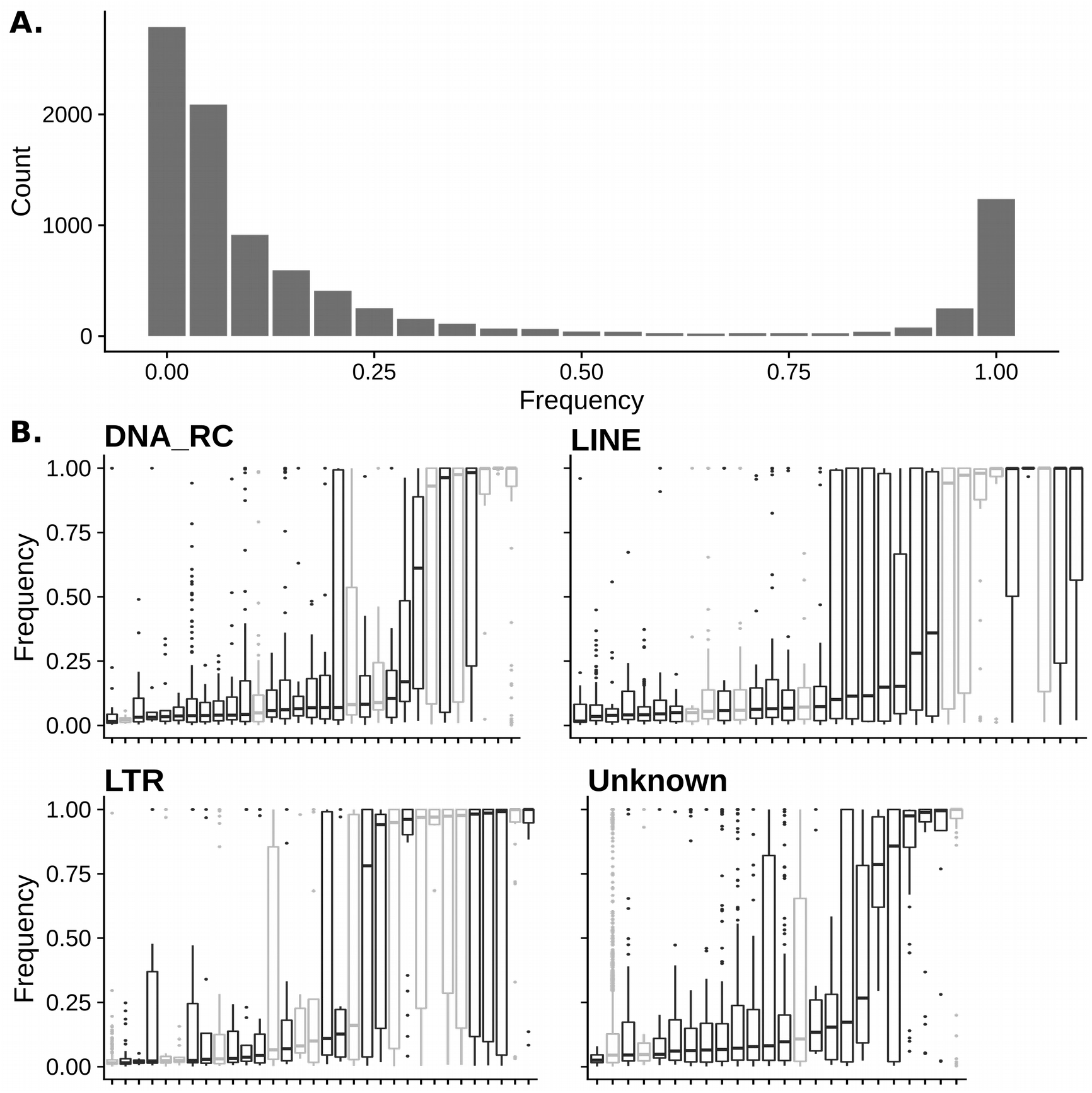
TE activity in the *D. suzukii* reference population from Watsonville (USA). A. Frequency distributions of TE insertions. B. Population frequencies for each TE family (in black) or pseudofamily (in gray). Only families/pseudofamilies with more than 10 insertions in the reference population are shown. DNA and Rolling Circles (RC) have been grouped for graphical reasons.

### Demography as driver of TE contents in *D. suzukii* populations

Our estimation of TE abundance in the 22 genotyped *D. suzukii* populations (fig. 3*A*) indicates substantial variation across populations, with significantly more TEs in invasive than in native populations and a strong correlation with the Watterson estimate of genetic diversity obtained from SNPs corresponding to a proxy of population effective size (fig. 3*B, C*). The mean number of insertions per Haploid Genome (HG) and per population was 2,793, ranging from 2,113 in the Chinese population CN-Nin to 3,129 in the Hawaiian population (US-Haw). There was a significant effect of the continent on the mean number of families/pseudofamilies per population: American and European populations had more families/pseudofamilies than native populations (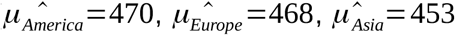, Kruskal-Wallis chi-squared = 10.505, df = 2, p-value = 0.0052). American and European populations also had more insertions per HG than native populations (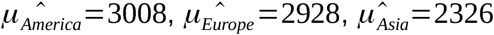, Kruskal-Wallis χ^2^ = 14.4, df = 2, p-value = 7.3×10^−4^). We found a negative linear correlation between the total number of insertions per HG and per population and the Watterson estimate of genetic diversity obtained from SNPs 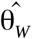, a proxy of population effective size (t = -13.415, df = 20, p-value = 1.8×10^−11^, fig. 3*C*). The variation of 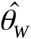 explains a large proportion of the variance in the total number of insertions per HG across the populations (R^2^ = 0.90). The correlation remains significant when considering only native populations (t = -5.22, df = 4, p-value = 6.4×10^−3^), or only invasive populations (t = - 3.06, df = 14, p-value = 8.6×10^−3^), or only European populations (t = -5.46, df = 6, p-value = 1.6×10^−3^), but not when considering only American populations (t = -1.89, df = 6, p-value = 0.11). The correlation between the number of insertions per HG per population and 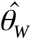 was also assessed individually for the 83 TE families/pseudofamilies showing an amplitude of variation superior or equal to 3 copies per HG. After a Benjamini-Hochberg correction for multiple testing, we found a significant correlation for 63 TE families (p-adjusted < 0.05).

**Figure 3:**
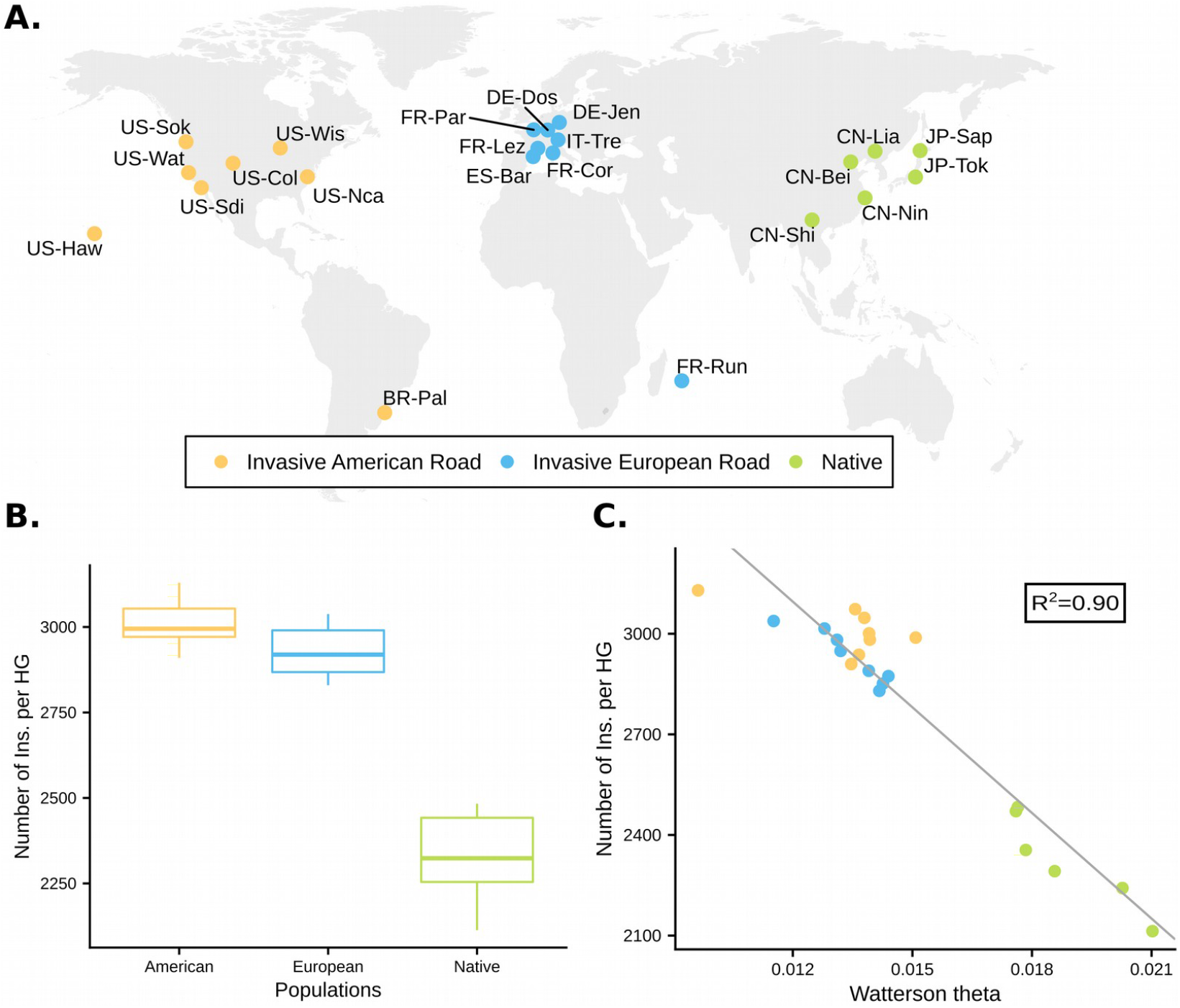
TE dynamics in native and invasive *D. suzukii* populations. A. Geographic location and historical status of the 22 *D. suzukii* population samples genotyped using a pool-sequencing methodology. Population samples from the native range are in green and those from the invaded range are in orange (American invasion route) or blue (European invasion route) (Fraimout et al. 2017). B. TE content in *D. suzukii* populations, as the numbers of insertions per haploid genome (HG). C. Correlation between TE content and Watterson’s theta in *D. suzukii* population samples.

### Environmental and genotypic effects on TE abundance

Because 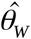 did not explain all the observed variation in TE abundance among the 22 sampled populations, we tested the effect of two other factors: the environmentally induced changes in TE activity and the genetically determined changes.

To test for an effect of environmentally induced changes in TE activity, we used Partial Mantel tests. We tested the correlation between 19 bioclimatic variables and TE family abundance, for the 83 TE families showing an amplitude of variation superior or equal to 3 copies per HG, correcting for population structure. After correction for multiple testing we did not find any significant correlation (Benjamini-Hochberg correction for multiple testing, p-adjusted < 0.05).

To evaluate the effect of genetic variation on TE abundance we performed a genome-wide scan for association using methods controlling for population structure. To that end we relied on the 13,530,656 bi-allelic variants (mostly SNPs) previously described on the same data set (Olazcuaga et al. 2020) and searched for association with the population abundance of the 83 TE families/pseudofamilies mentioned above using the BayPass software. Globally, we found 4,856 genomic regions showing evidence of association with population abundance of at least one TE family. Each region spanned at least 1 kb on the reference assembly and included one or several significant SNP/InDel separated by less than 1 kb (significance threshold: Bayes Factor (BF) >20). On average each region was associated with the number of insertions per HG of 1.37 families (min=1, max=69) and contained 2.40 SNPs/InDels (min=1, max=49). 306 (6.30%) regions overlapped with repeated sequences as annotated in the reference genome, which is less than expected by drawing SNP/InDel associated regions randomly (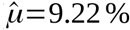, q_0.025_=7.60%, q_0.975_=11.16%; supplementary fig. S4*A*). Only 14 of these regions contain a TE of the same family/pseudofamily as the TE abundance they were associated with. Regarding genes, 2,843 (58.55%) regions were associated with at least one gene, which is less than expected under random expectations (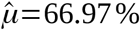, quantile_0.025_=62.40%, quantile_0.975_=70.76%; supplementary fig. S4*B*). Due to their known role in the activity of TEs, we further searched for enrichment in genes encoding transcription factors and piRNA pathway effectors among the genes located within our candidate regions. We did not observe any significant enrichment in genes encoding transcription factors (Observed: 13.63%, Expected: 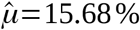, quantile_0.025_=12.00%, quantile_0.975_=20.60%; supplementary fig. S4*C*) nor in genes involved in the piRNA pathway (Observed: 0.33%, Expected: 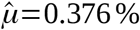, q_0.025_=0.00%, q_0.975_=1.18%; supplementary fig. S4*D*). Among the top 10 regions, corresponding to the regions associated with the highest number of TE families/pseudofamilies, two appeared to be non genic, four could not be attributed to *D. melanogaster* genome, three were associated with the mitochondrial genome and one was associated with *blot*, a member of the sodium- and chloride-dependent neurotransmitter symporter family (https://flybase.org/reports/FBgn0027660).

### A small number of putatively adaptive TE insertions

We investigated the presence of putatively adaptive insertions using a genome scan combining three methods controlling for population structure implemented in BayPass (Olazcuaga et al. 2020). First, we assessed overall differentiation (based on the XtX statistics). Second, we studied allele frequencies differences between two groups of populations (based on the C_2_ statistics): American invasive *vs* native populations (C_2_^Am^), European invasive *vs* native populations (C_2_^Eu^), all invasive *vs* native populations (C_2_^WW^). Third, we carried out genome-wide association with each of the 19 bioclimatic variables (based on the BF).

The genome scan was conducted on 7,004 polymorphic TE insertions (MAF > 0.025, 5,944 autosomal insertions and 1,060 X-linked insertions treated separately). We identified a total of 15 putatively adaptive insertions (13 located on autosomal and three on X-linked contigs) (table 1; fig. 4). Nine of these insertions were outliers when considering the global differentiation statistics XtX. Note that their frequencies were distinct between native Chinese (low frequencies) and native Japanese populations (high frequencies). One insertion was an outlier for both the XtX and C_2_^Am^ statistics. Finally, the last five insertions were outliers for the C_2_^WW^ statistics. No significant association was found between TE insertion frequencies and the 19 bioclimatic variables investigated.

**Table 1:** Description of the 15 putatively adaptive TE insertions. Each insertion is an outlier when considering one or a combination of the global differentiation statistics (XtX) and statistics contrasting allelic frequencies between native populations and populations of the invasive American road (C_2_^Am^) or populations of the invasive European road (C_2_^Eu^) or all invasive populations (C_2_^WW^).

| Insertion | Statistics | Gene vicinity | Outlier SNP nearby | A/X | TE Order |
| --- | --- | --- | --- | --- | --- |
| 1 | $C_2^{Am} - XtX$ | ASPP | F | A | Unknown |
| 2 | $C_2^{WW}$ | dia | F | A | Unknown |
| 3 | $C_2^{WW}$ | - | T | A | DNA |
| 4 | $C_2^{WW}$ | NA | F | X | Unknown |
| 5 | $C_2^{WW}$ | inaE | F | X | Unknown |
| 6 | $C_2^{WW}$ | - | F | X | DNA |
| 7 | $XtX$ | Mical | F | A | DNA |
| 8 | $XtX$ | CG30015 | F | A | Unknown |
| 9 | $XtX$ | - | F | A | Unknown |
| 10 | $XtX$ | CR31386 | F | A | Unknown |
| 11 | $XtX$ | - | F | A | Unknown |
| 12 | $XtX$ | Dop1R2 | F | A | Unknown |
| 13 | $XtX$ | jing | F | A | Unknown |
| 14 | $XtX$ | CG14282 | F | A | Unknown |
| 15 | $XtX$ | GATAe | F | A | Unknown |

Note.—The fourth column indicates whether a SNP potentially evolving under positive selection had been detected less than 5 kb away in Olazcuaga et al. (2020) (F=False, T=True). The fifth column indicates whether the insertion is located on an autosomal (A) or X-linked contig (X).

**Figure 4:**
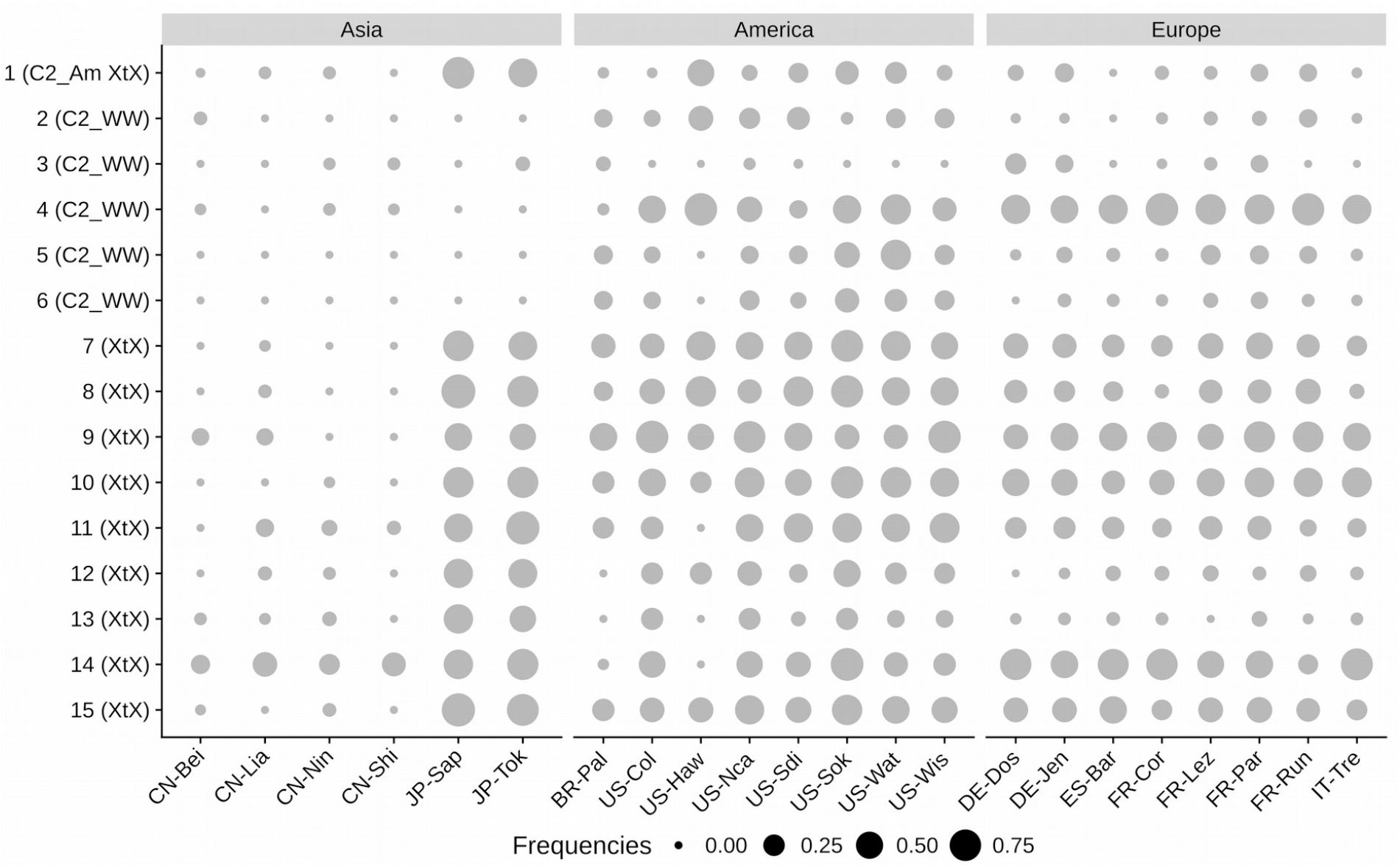
Frequencies of each of the 15 putatively adaptive insertions in the 22 *D. suzukii* populations. Insertion number is indicated on the left together with the associated BayPass statistics. XtX corresponds to a global differentiation statistic, C_2_ to a statistic contrasting allelic frequencies between native populations and populations of the invasive American road (C_2_^Am^) or populations of the invasive European road (C_2_^Eu^) or all invasive populations (C_2_^WW^_)_).

One of the 15 putatively adaptive insertions was close (*i*.*e*., 399 bp away) to a SNP/InDel that had previously been identified in a region potentially associated with *D. suzukii* invasive success (table 1) (Olazcuaga et al. 2020). For one insertion we did not find any homologous regions in *D. melanogaster*, four others were in genomic regions without any genes, and the ten remaining were associated with genes.

We further investigated signatures of selection around candidate insertions by estimating local Tajima’s D statistics in the SNP/InDel dataset. Low values of Tajima’s D indicate an excess of rare mutations, one possible signature of a selective sweep due to positive selection. To test if each of our candidate insertions were associated with selective sweeps, we computed the linear correlation between its frequency and local Tajima’s D values (supplementary fig.S5). Five statistically significant correlations were found corresponding to the insertions n°4, 9, 10, 12 and 15 (Pearson’s product-moment correlation, p < 0.05). Only a single insertion was associated with an extreme local Tajima’s D (insertion n°15; Tajima’s D < quantile_0.05_), and only for a single population. The visualization of Tajima’s D at a larger scale (*i*.*e*., 10 kb upstream - 10 kb downstream the insertion) confirms the lack of strong effect of the investigated insertions on Tajima’s D (supplementary fig. S6). It is worth noting that, if the effect of our candidate TE insertion on Tajima’s D is globally low, a close investigation of Tajima’s D suggests that, at least in some cases, it is the absence rather than the presence of the insertion that may be adaptive. As a matter of fact, while the correlation implying an extreme local Tajima’s D was negative, the four other significant correlations between local Tajima’s D and insertion frequency were positive.

## Discussion

For most species the repeatome is still a poorly known genomic compartment and much remains to be understood regarding its variability, dynamics, functional and fitness impacts. This is all the more important given that TEs appear to be ubiquitous, prompt to invade new genomes (Kofler, Hill, et al. 2015), and they may drastically impact the host phenotype (Nikitin and Woodruff 1995; Daborn et al. 2002; Van’t Hof et al. 2016). Here we capitalized on a recently generated long-reads genome assembly and a large set of populational PoolSeq data (Olazcuaga et al. 2020; Paris et al. 2020) to thoroughly portray the TE content of the non model invasive species *Drosophila suzukii*.

### An abundant, unevenly distributed and active repeatome

The observed 47% of TEs in the genome of *D. suzukii* confirmed the outlier position of this species within the Drosophila genus regarding the global amount of TEs. Our estimate is somewhat higher than those reported in previous studies in *D. suzukii* (Chiu et al. 2013; Ometto et al. 2013; Sessegolo Camille et al. 2016; Paris et al. 2020). Considering that the assembly of repeats is often impossible using short paired-end (PE) reads (Rius et al. 2016), it is not surprising that we recovered more TEs in a long reads genomic assembly than previous studies investigating TE contents using PE reads assemblies (Chiu et al. 2013; Ometto et al. 2013). In addition, we here performed a *de novo* reconstruction of TE sequences, which allowed us to identify more TE families/pseudofamilies, as compared to the previous research work based on the same assembly (35 %) (Paris et al. 2020). Overall, *de novo* reconstruction of TE sequences from long read assemblies, such as the 15 Drosophila species assemblies recently generated using nanopore sequencing (Miller et al. 2018), should greatly improve our knowledge of TE diversity in Drosophila.

In agreement with the gene disruption hypothesis and observations in a variety of species (Bartolomé et al. 2002; Medstrand et al. 2002; Wright et al. 2003), we observed a depletion of TE copies in gene-rich regions of the *D. suzukii* genome. Although it is likely that TEs are strongly selected against in these regions due to their negative effect on gene function or expression (Lee and Karpen 2017; Mérel et al. 2020), it is also possible that TE copies are depleted in these regions because they promote ectopic recombination. In agreement with the latter hypothesis, gene-rich regions are also known to display high recombination rate in *D. melanogaster (Adams et al. 2000)*. The generation of a genomic map of recombination rates in *D. suzukii* would be needed to disentangle the respective effects of ectopic recombination and gene disruption.

At the chromosomal scale, we did not find a lower density of TEs on the X chromosome compared to autosomes. This pattern indicates that, if X-linked recessive insertions are more efficiently selected against than autosomal insertions, the effect on TE abundance is either low or balanced by another process. When comparing only gene-rich regions, we even found a higher density of TEs on the X chromosome than on autosomes. Three non-mutually exclusive explanations can be invoked: (i) there may be a higher insertion rate on the X chromosome, similar to what was previously found in *D. melanogaster* (Adrion et al. 2017); (ii) the recombination rate may be lower on the X chromosome, and thus a stronger Muller’s ratchet; and (iii) the strength of selection may be reduced by a smaller effective population size for the X chromosome.

Similarly to what has been found in *D. melanogaster* and *D. simulans* (Kofler, Nolte, et al. 2015) and to what is probably common among Drosophila species (Hill 2019), the pattern of TE insertion frequencies in *D. suzukii* is compatible with an active repeatome. We found differences in the mean insertion frequency between TE orders, which suggests differences in activity but could also result from variation in the strength of purifying selection acting against the different orders (Petrov et al. 2003; Lee and Karpen 2017) Considering the trap model of TE dynamics (*i*.*e*. a model in which newly invading TEs are quickly inactivated by host defense (Zanni et al. 2013; Kofler et al. 2018)), an active repeatome suggests a recurrent turnover of TEs, potentially due to horizontal transfer events. Investigating TE activity in *D. melanogaster* and *D. simulans*, Kofler and colleagues (Kofler, Nolte, et al. 2015) suggested that such a turnover is influenced by the colonization history of those species. They propose that the high activity of DNA transposons in *D. simulans* results from horizontal transfer events from *D. melanogaster* during *D. simulans* worldwide colonization. In agreement, we detected more families/pseudofamilies in invasive populations of *D. suzukii* than in the native ones, suggesting that new TE families may have been acquired during the recent colonization of new areas. However, because the TE database used here relies on a reference genome obtained from individuals originating from America (*i*.*e*. from the Watsonville population), one may expect to find much more families/pseudofamilies in American than European populations. Yet this is not what we observed. This could be due to admixture between American and European populations. However, population genetics studies have shown that gene flow between the two continents is limited if not absent (Fraimout et al. 2017). It is thus possible that, for technical reasons, we are simply missing some families that are less abundant in the Asian native range of the species. The comparison of long read assemblies of genomes generated from individuals originating from the three continents (Asia, America and Europe) should help shedding light on this issue.

### Demography, rather than environment or genotype, drives TE content

In agreement with the Lynch and Connery hypothesis (Lynch and Conery 2003), we found that the TE content in *D. suzukii* is negatively correlated with the Watterson estimate of genetic diversity 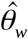 which may be viewed as a proxy of the population effective size N_e_. The negative correlation between 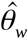 and TE content was significant when considering only European invasive populations, invasive populations as a whole, or only native populations, but was not significant when considering only American invasive populations. Although a few studies suggest an increase of TE content following colonization (Nardon et al. 2005; García Guerreiro et al. 2008; García Guerreiro and Fontdevila 2011; Talla et al. 2017), to our knowledge it is the first time that a correlation between TE content and N_e_ is found at the intraspecific level. Although several factors may affect N_e_, the variation observed is likely to result from demographic processes. Indeed, both European and American invasive populations have encountered bottlenecks (Fraimout et al. 2017). In agreement with this idea, the invasive population from Hawaii, which experienced the strongest bottleneck (Fraimout et al. 2017), showed the smallest 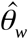 values. It is interesting to note that the negative correlation between 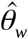 and TE content remains significant when considering only native populations suggesting that other demographic event than bottleneck may also be involved (e.g. different stable effective population sizes and gene flow patterns),

Our analysis is controlled for sequencing bias, *i*.*e*. coverage and insert size, and we are confident in the biological significance of the correlation observed here. However, it is worth stressing that our dataset of TE insertions corresponds to a small fraction of the repeatome. Indeed, the mean number of insertions per HG per population is markedly below the number of TE copies recovered in the reference genome. We believe that this is due to an impossibility to properly call TE insertions when TEs are too close or even nested (Vendrell-Mir et al. 2019). It is thus possible that the negative correlation that we found here exists only for some part of the genome. Especially it is likely that regions of low TE density, where most of TE insertions are polymorphic, display the strongest answer to a reduction of selection efficacy. This is simply because polymorphic insertions can increase in frequency while fixed insertions cannot. One could also argue that the efficiency of selection is a function of the product between N_e_ and s (with s the selection coefficient). Therefore, the effects of a reduction of Ne should be especially marked in regions where selection against TEs is strong, such as TE-poor / gene-rich regions.

We found no significant effect on TE abundance for all the 19 environment variables tested. This might be surprising at first sight given the large number of studies showing an association between TE activity and external factors, such as temperature or viral infection (García Guerreiro 2012; Ryan et al. 2016; Horváth et al. 2017; Roy et al. 2020) Several factors may explain this discrepancy. First, it is important to notice that, in Drosophila, most of these studies rely on lab experiments, some of them exploring environmental conditions unlikely *in natura* (see (García Guerreiro 2012) for a review). To our knowledge none of these studies established a link between TE activity and natural environment without any possible confounding effect from population structure and demographic features. Second, as often in Drosophila, most of such research works were carried out on the same particular species,*D. melanogaster*, so that so far we do not know much about interspecific variability. Third, although partial Mantel tests allowed revealing 15 significant correlations between TE abundance and environmental variables in *A. thaliana* populations (Quadrana et al. 2016), we consider our results as conservative, especially regarding the long discussion about the statistical performance of partial Mantel tests (Diniz-Filho et al. 2013). More sophisticated statistical methods may be needed to tackle such relationships into more details.

Considering that several studies on Drosophila suggest a genotype effect on TE activity (Biémont et al. 1987; Pasyukova and Nuzhdin 1993; Díaz-González et al. 2011; Adrion et al. 2017), we performed a GWAS on TE abundance to assess this effect in natural populations and identify the genomic regions involved. Overall, we found *ca*. 5,000 genomic regions associated with TE abundance. These regions were not enriched in transcription factor genes nor genes of the piRNA pathway. As far as we know, no such GWAS study has been carried out in Drosophila populations. Our results are somewhat similar to those found in *A. thaliana*, in which although a strong causal link between one transcription factor and the abundance of two TE families was found, no enrichment for any particular function was observed (Quadrana et al. 2016). Comparative genomics between closely related species may help identify a general pattern. Especially, one could lead the same study using available *D. melanogaster* PoolSeq data (Kapun et al. 2020), and focus on genes identified in both *D. melanogaster* and *D. suzukii*, as they might be likely to play a key role in the modulation of TE activity.

### A potential adaptive role for a limited number of TEs

Similar to studies investigating TE adaptive potential in *D. melanogaster* populations (González et al. 2008; González et al. 2010; Rech et al. 2019), we found several putatively adaptive TE insertions in our *D. suzukii* dataset. Overall, we found 15 insertions, six of which likely to have eased the worldwide invasion of *D. suzukii*. It is important to note that we are probably missing some insertions, and thus likely underestimating the number of adaptive insertions sites.

Overall, we did not capture a strong signal of a selective sweep near the candidate adaptive TE insertions. This may be due to overall large effective population sizes as suggested in (Olazcuaga et al. 2020), but also to the fact that Tajima’s D is unlikely to detect soft selective sweep, *i*.*e*. adaptation from standing variation or multiple successive beneficial mutations (Pennings and Hermisson 2006). An appealing perspective would be to sequence candidate regions in individual strains and use a haplotype-based analysis. For example, the recently introduced Comparative Haplotype Identity (xMD) statistics (Lange and Pool 2016; Villanueva-Cañas et al. 2017) has been shown to perform well for soft sweeps. If the effect of our candidate TE insertion on Tajima’s D is globally low, it highlighted the possibility that the absence rather than the presence of the insertion may be adaptive, at least for some of our candidate insertions. More specifically, for four insertions a positive correlation was found between local Tajima’s D and insertion frequency. However, the only extreme local Tajima’s D was found in the population where the putatively adaptive insertion is at its highest frequency, indicating that it is probably the insertion itself rather than the absence that might be adaptive.

One added value to our analysis based on GWAS is that the same type of analysis has been carried out using SNPs/InDel (Olazcuaga et al. 2020). The authors of this study found 204 markers strongly associated with invasion success distributed over the whole genome. If we compare this number to our six TE insertions, it seems unlikely that TEs solely may explain the genetic paradox of invasive species (Stapley et al. 2015). It is worth noting that the level of variation remains high in invasive *D. suzukii* populations (Fraimout et al. 2017). Hence, it would be interesting to carry out similar analyses in invasive species that experienced a more intense depletion of genetic variation during invasion (Prentis et al. 2009; Zhang et al. 2010; Roux et al. 2011) to assess whether TEs are more likely to be adaptive in invasive populations with low levels of genetic diversity.

At first sight our finding of 15 putatively adaptive polymorphic insertions in worldwide populations of *D. suzukii* contrasts with the 41 to 300 putatively adaptive polymorphic insertions found in worldwide populations of *D. melanogaster* (Rech et al. 2019). The difference is even more blatant considering that we analyzed 7,004 polymorphic insertions, against ∼800 in (Rech et al. 2019). This suggests a largely higher rate of TE induced adaptations during *D. melanogaster* invasion and this despite the much larger, still active and diverse repeatome of *D. suzukii*. This discrepancy could have several non-exclusive explanations. First, it may be due to historical differences between the two species. *D. melanogaster* experienced a relatively slow and ancient worldwide invasion that started from Africa about ∼15,000 ya, whereas *D. suzukii* came out from its native range in Asia only a few decades ago (Stephan and Li 2007; Fraimout et al. 2017). Second, the discrepancy may result from intrinsic species differences with respect to the repeatome contents. For example, *D. melanogaster* TEs could possess more environment responsive sequences that might be co-opted by the host. Third, it may be due to differences in the methodology used for the two species. Our analysis relies essentially on the research of overly differentiated TEs across populations with a correction for population structure (Gautier 2015; Olazcuaga et al. 2020), whereas in the analysis used for *D. melanogaster* there is no direct methodological control for population structure. In the *D. melanogaster* study (Rech et al. 2019), a TE insertion is considered as putatively adaptive if it is present at high population frequency (from 10% to 95%), and is located in genomic regions where recombination rate -and so selection efficacy - is high (*ca*. 300 putatively adaptive insertions). Further evidence is collected using a combination of three haplotype-based tests to detect selective sweeps in the vicinity of candidates, and statistical treatments based on Fst estimations (with 84 insertions confirmed by at least one test). Applying our statistical methodologies to the *D. melanogaster* dataset, which also consist in PoolSeq data, would help to determine if methodology differences can explain the observed discrepancy. Finally, one could ultimately rely on experimental evolution, applying the same selective pressure to different Drosophila species, to test for an impact of intrinsic species differences on TE adaptive potential.

Our study of TE induced adaptation strongly calls for a validation of candidate insertions. Allele specific expression assays would allow evaluating if these insertions affect nearby gene expression (Gonzalez et al. 2009). This would consist in testing a difference of nearby gene expression between the two alleles of an F1 hybrid between strains with and without the insertion. While such test should control for genotype effect, as compared to a simple test of differential expression between strains, it does not preclude for an effect of a SNP/InDel close to the insertion. Using a CRISPR/Cas9 methodology would also allow (in)validate that the TE(s) of interest is the causative agent of gene expression change and would allow direct testing for a phenotypic effect.

### Conclusion

Our study illustrates the value of an approach combining a long reads based genome assembly, a *de novo* reconstruction of TE sequences, and PoolSeq population data, to characterize the repeatome of a non model species. Our set of analyses especially highlighted that the particularly large *D. suzukii* repeatome is probably active and shaped by purifying selection, similar to that of *D. melanogaster*’s. Additional data, such as local recombination rate, would also help us shed light on the nature of selection acting on TEs. The analysis of TE abundance variations in invasive and native populations suggests that a reduction of purifying selection intensity, in response to demographic processes, can significantly increase TE content. Our study also indicates that positive selection may act on TE insertions in response to selective factors that remains to be determined. Experimental validation will allow to (in)validate a functional impact of our putatively adaptive insertions. Overall, the natural extent of the trends we uncovered here should be explored into more details, for instance through the application of similar methods to other (invasive) species that would allow to evaluate the impact of a stronger bottleneck on both TE content increase and TE adaptive potential.

## Materials & Methods

### Creation of a TE database

A TE database was created by merging previously established consensus of Drosophila TE families and *de novo* reconstructed consensus of *D. suzukii* TE families. The previously established consensus were obtained by extracting all Drosophila consensus annotated as DNA, LINE, LTR, Other, RC, SINE and Unknown from Dfam and Repbase databases (release 2016-2018 for both) (Hubley et al. 2016; https://www.girinst.org/repbase/). Full LTR element sequences were reconstructed by merging LTRs and their internal parts. *De novo* reconstruction was performed using an assembly of an American strain from Watsonville, sequenced using PacBio long reads technology, and the REPET package (v2.5) (Flutre et al. 2011; Paris et al. 2020). Unless otherwise specified, the options were used as in the default configuration file. Briefly, the genome assembly was cut into batches and aligned to itself using blastn (ncbi-blast v2.2.6) (Altschul et al. 1990). High-scoring Segment Pairs (HSPs) were clustered using Recon (v1.08) and Piler (v1.0) (Bao and Eddy 2002; Edgar and Myers 2005). A structural detection step was performed using LTRHarvest from the GenomeTools package (v1.5.8) (Ellinghaus et al. 2008; Gremme et al. 2013). LTRHarvest-produced sequences were clustered using blastclust. Consensus sequences were created for each cluster using MAP (Huang 1994). Additional consensus sequences were generated using RepeatScout (v1.0.5) (Price et al. 2005). All consensus, *i*.*e*. from Recon, Piler, LTRHarvest and RepeatScout, were further submitted to a filtering step. Sequences were retained only if they produced at least 3 hits against the genome assembly with at least 98% query coverage (blastn, blast 2.6.0+). Structural and coding features were identified and used to classify consensus (see Hoede et al. (2014) for classification details, the used libraries were ProfilesBankForREPET_Pfam27.0_GypsyDB.hmm, repbase20.05_aaSeq_cleaned_TE.fsa, repbase20.05_ntSeq_cleaned_TE.fsa). Single satellite repeats, potential host genes and unclassified sequences were filtered out. Since REPET can easily mis-annotate any pair of repeats separated by a spacer as TRIM or LARD, those sequences were also removed (Arkhipova 2017). Remaining sequences were further annotated by homology to previously established consensus of Drosophila TE families. Homology was determined using RepeatMasker (-cutoff 250, v 1.332) (http://www.repeatmasker.org/). We followed the rules below: 1) if all hits belonged to the same superfamily, the sequence was annotated as corresponding to that particular superfamily and order; 2) if hits from different superfamilies were observed the sequence was considered as ambiguous; 3) without any hit, the sequence was annotated as unknown. Ambiguous sequences were manually curated, sequences which could be unambiguously attributed to one superfamily according to hits and proteic domains were kept (proteic domains were investigated using NCBI Conserved Domain Search (https://www.ncbi.nlm.nih.gov/Structure/cdd/wrpsb.cgi). Finally, consensus were clustered in families using UClust (-id 0.80, -strand both, –maxaccepts 0 –maxrejects 0; v11.0.667) (Edgar 2010). The annotation, superfamily and order, attributed to each cluster, *i*.*e*. each family, is the annotation of the longest sequence in the cluster. The generated TE database is accessible at: https://github.com/vmerel/Dsu-TE.

### Annotation of the reference genome

To recover TE fragments and TE genomic sequence occupancy, the reference genome assembly was masked using RepeatMasker and the above TE database (-gccalc, -s, -a, -cutoff 200, -no_is, -nolow, -norna, -u; v 1.332) (http://www.repeatmasker.org/). TE density was evaluated as the number of TE fragments completely within non overlapping genomic windows of 200 kb. TE copies were reconstructed from TE fragments using OneCodeToFindThemAll (Bailly-Bechet et al. 2014). Gene density was computed from a run of augustus (–species=fly, – strand=both, –genemodel=complete; v2.5.5) (Stanke et al. 2008) as the number of genes completely within non overlapping genomic windows of 200 kb. Promer was used to generate alignments between *D. melanogaster* and *D. suzukii* assemblies and establish syntenic relationships (MUMmer v3.23) (Kurtz et al. 2004). *D. melanogaster* masked assembly was downloaded from UCSC Genome Browser (dm6; http://hgdownload.soe.ucsc.edu/goldenPath/dm6/bigZips/). *D. suzukii* masked assembly was retrieved from RepeatMasker output (see above). The promer output was filtered out using the delta-filter module in order to obtain a one-to-one mapping of reference to query (-q, -r). A file containing alignment coordinates for alignments of minimum length 100 bp, and in which overlapping alignments were merged, was generated with the show-coords module (-b, -L 100, - r). Because the abundance of repeated sequences and the use of masked assemblies may result in multiple small alignments, alignments separated by less than 20 kb were merged using a custom script. Note that only alignments implying the 2L, 2R, 3L, 3R, X and 4 chromosomes of *D. melanogaster* were kept at this step and if a *D. suzukii* contig aligned to several *D. melanogaster* chromosomes only the best pair was conserved (*i*.*e*. the pair producing the longest alignment). A graphical visualization of the results was produced using Circos (Krzywinski et al. 2009).

### Fly samples and pool sequencing

Pool-sequencing (PoolSeq) data originate from Olazcuaga et al. (2020) where the detailed associated protocol is described. Briefly, adult wild flies were sampled between 2013 and 2016 from 22 localities of both native and invasive areas (fig. 3*A*) (Fraimout et al. 2017). Six samples were collected in the native Asian area, more precisely in four Chinese and two Japanese localities. The remaining 16 samples were chosen to be representative of two separate invasion roads: the American invasion road and the European invasion road. The American invasion road is represented by one Hawaiian sample, one Brazilian sample and six samples from the United States. The European invasion road corresponds to two German samples, four French samples (including one from La Réunion Island), one Italian sample and one Spanish sample. For each population sample, DNA extraction was performed from the thoraxes of 50 to 100 flies and used to prepare paired-end (PE) libraries (insert size of ∼550 bp). PE sequencing was achieved using a HiSeq 2500 from Illumina to obtain 2×125 bp reads. Reads were trimmed using the trim-fastq.pl script in the PoPoolation package (–min-length 75, –quality-threshold 20; v1.2.2) (Kofler et al. 2011).

### TE frequency pipeline

To obtain TE insertion frequencies in PoolSeq samples a calling of TEs was done using PoPoolationTE2 (Kofler et al. 2016), the reference genome and the newly constructed database. To make sure that no reads from TE sequences could map on the masked assembly, TE reads were simulated, mapped on the masked assembly and aligned positions were also masked. Reads simulation was performed using the script create-reads-for-te-sequences.py (Kofler et al. 2016): reads of 125 bp reads, coverage of 1024 X per TE sequence in the database. Because we do not expect a split read based TE calling tool such as PoPoolationTE2 to accurately call for insertions shorter than the insert size, TE sequences shorter than 500 bp were removed before calling. Moreover, as PoPoolationTE2 filters out insertions with reads mapping on more than one family, families with cross-mapping were grouped in pseudofamilies. Two families were brought together if at least 1% of reads from one sequence of the first family were mapped on a sequence of the second family (read simulation: 125 bp reads, coverage of 100 X per consensus). Concerning the TE calling, reads were mapped using bwa bwasw (v0.7.17) (Li and Durbin 2010) and paired-end information restored using the se2pe script provided with the PoPoolationTE2 package (v1.10.04) (Kofler et al. 2016). One unique ppileup file was generated with all samples specifying a minimum mapping quality of 15. The remaining modules of PoPoolationTE2 were used as follow: identifySignatures: –mode joint, –signature-window minimumSampleMedian, –min-valley minimumSampleMedian, –min-count 2; updatestrand: –map-qual 15, –max-disagreement 0.5; frequency; filterSignatures –min-coverage 10, –max-otherte-count 2, –max-structvar-count 2; pairupSignatures –min-distance - 200, –max-distance 300. The final output contained frequencies in the 22 populations for each called TE insertion. See supplementary methods for the validation work on simulated data.

### TE abundance pipeline

TE abundances, as the numbers of insertions per HG per population, were estimated in PoolSeq samples by summing insertion frequencies in each sample. Since this pipeline also relies on the estimation of TE frequencies in PoolSeq samples, it is very similar to the TE frequency pipeline. However, the last steps were modified to account for differences in coverage and insert sizes between samples and to allow an unbiased comparison of TE abundance across samples. After the ppileup step the following analyses were performed: subsamplePpileup: –target-coverage 30; identifySignatures –mode separate, –signature-window minimumSampleMedian, –min-valley minimumSampleMedian, –min-count 2; updatestrand: –map-qual 15, –max-disagreement 0.5; frequency; filterSignatures: –min-coverage 10; –max-otherte-count 2; –max-structvar-count 2; pairupSignatures: –min-distance - 200; –max-distance 300. See supplementary methods for the validation work on simulated data.

### Evaluation of population genetics statistics

We estimated Watterson’s theta 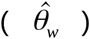 and Tajima’s D statistics in non-overlapping 1000 bp windows using PoPoolation (v1.2.2) (Kofler et al. 2011). Forward and Reverse trimmed reads were mapped separately using bwa aln (-o 2 -d 12 -e 12 -n 0.01; v0.7.17) (Li and Durbin 2010). A paired-end alignment file was generated using bwa sampe. Reads were filtered for a minimum mapping quality of 20 and a pileup file generated with samtools (v1.7) (Li et al. 2009). Each pileup file was split into two files: one corresponding to autosomal contigs and another corresponding to X-linked contigs (autosomal and X-linked contigs as determined in Olazcuaga et al. (2020)). PoPoolation was used as follows: –min-count 2 –min-coverage 8 –max-coverage 250 –min-qual 20. The pool-size argument was modified accordingly between autosomal and X-linked pileup.

### Genome Wide Association Study with TE family abundance

All genome scans were performed using BayPass (v2.2) (Gautier 2015; Olazcuaga et al. 2020), a package aiming at identifying markers evolving under selection and/or associated to population-specific covariates, taking into account the shared history of the populations. For each SNP/InDel previously called in these PoolSeq samples (Olazcuaga et al. 2020), we estimated 83 Bayes Factors (BF), reflecting their association with the number of insertions per HG of 83 families/pseudofamilies (based on a linear regression model). The 83 chosen TE families/pseudofamilies were those displaying an amplitude of variation of at least three insertions per HG across the complete dataset. To improve computing time BayPass was run on data subsets. Data concerning TE abundance was split into three subsets of 28, 28 and 27 families, respectively. For SNPs/InDel, we used the data subsets of Olazcuaga et al. (2020), for which the 11,564,472 autosomal variants are divided into 154 subsets and the 1,966,184 X-linked variants into 26 subsets. Since we used the importance sampling algorithm implemented in Baypass to assess BFs, and single run estimations may be unstable, a total of three runs were performed for each combination of TE subsets-SNP/InDel subsets and the median of BFs computed (Gautier et al. 2018). Note that different pool size files were used for autosomal and X-linked variants to take into account differences in the number of autosomes and X chromosomes in each PoolSeq sample. In accordance to Jeffrey’s rule, a SNP/InDel was considered as associated with a TE family/pseudofamily abundance for a BF superior to 20 deciban (dB) (Jeffreys 1961).

SNP/InDel locations were used to define genomic regions associated with TE abundance. Variants were gathered if separated by less than 1 kb. If the spanned genomic interval was less than 1 kb or if a variant could not be found, the region was obtained by adding 500 bp on both sides. For each region we looked for overlapping TEs using the RepeatMasker annotation (gff file, see *Annotation of the reference genome*). We also investigated gene content. First, we retrieved homologous regions in the *D. melanogaster* genome using BLAT against the *D. melanogaster* masked assembly downloaded from UCSC Genome Browser (http://hgdownload.soe.ucsc.edu/goldenPath/dm6/bigZips/; BLAT v.36×4, -t=dnax -q=dnax). We then checked for genes overlapping the best hit subject sequence using the UCSC Genome Browser gff annotation file. Note that if the best hit score was lower than 100 we considered that no homologous region was retrieved. The number of transcription factor genes among the genes retrieved was obtained by comparing their IDs to those of the gene group Transcription factor on flybase (https://flybase.org/reports/FBgg0000745.html). Similarly, the number of genes involved in the piRNA pathway was obtained by comparing gene IDs to those listed in Ozata et al. (2019). In order to test if the candidate regions were enriched in TEs we generated random expectations by applying the above to 1000 randomly selected SNPs 250 times. For computing time reasons, for genes, transcription factor genes, or genes involved in the piRNA pathway, we used 500 randomly selected SNPs 125 times.

### Correlation between climatic variables and TE family abundance

Partial Mantel tests were used to test the correlation between bioclimatic variables and TE family abundance correcting for population structure (as in Quadrana et al. (2016)). 19 bioclimatic variables from the worldclim dataset (Fick and Hijmans 2017) were considered: annual mean temperature, mean diurnal range, isothermality, temperature seasonality, max temperature of warmest month, minimum temperature of coldest month, temperature annual range, mean temperature of wettest quarter, mean temperature of driest quarter, mean temperature of warmest quarter, mean temperature of coldest quarter, annual precipitation, precipitation of wettest month, precipitation of driest month, precipitation seasonality, precipitation of wettest quarter, precipitation of driest quarter, precipitation of warmest quarter, precipitation of coldest quarter. The 83 families with an amplitude of variation of at least three insertions per HG between populations were considered. The population structuring of genetic diversity is summarized by the scaled covariance matrix of population allele frequencies (Ω) estimated with Baypass, one autosomal subset randomly chosen was used (the correlation of the posterior means of the estimated Ω elements across SNP subsamples had previously been verified (Olazcuaga et al. 2020)). Partial Mantel tests were conducted using the R package ecodist (Goslee and Urban 2007). P-values were further adjusted to account for multiple testing applying the Benjamini-Hochberg correction (Benjamini and Hochberg 1995).

### Screening for putatively adaptive TE insertions

A genome scan for putatively adaptive TE insertions was performed using BayPass with the output of the TE frequency pipeline (v2.2) (Gautier 2015; Olazcuaga et al. 2020). Insertions with Minor Allelic Frequency (MAF) inferior to 0.025 were removed before the analysis. Autosomal and X-linked contigs were analyzed separately. Three statistics were computed to detect putatively adaptive TE insertions: XtX, C_2_ and the Bayes Factor (BF) for Environmental Association Analysis. Briefly, XtX corresponds to a global differentiation statistics, C_2_ contrasts allelic frequencies between user-defined groups of populations, and BF measures the support of the association between a marker and a covariate (usually an environmental variable). Because Bayes Factor was computed using the importance sampling algorithm, and single run estimations may be unstable, BF were estimated as the median over five estimates obtained from independent runs of Baypass (Gautier et al. 2018). In accordance to Jeffrey’s rule, a BF superior to 20 deciban (dB) was considered as decisive evidence supporting an association (Jeffreys 1961). XtX and C_2_ estimates came from one single run and simulation was used to determine a significance threshold. The R function simulate.baypass() provided within the BayPass package was used to simulate read count data (nsnp=10000, pi.maf=0). We used the physical coverage estimated from the ppileup file using the module stat-coverage of PoPoolationTE2 (Kofler et al. 2016). BayPass was run on this simulated dataset to estimate the null distribution of the XtX and the C_2_ statistics. An insertion was considered as overly differentiated (for XtX) or associated to the tested contrast (for C_2_) if the corresponding statistics exceeded the 99.9% quantile of the estimated null distribution. The populations whose frequencies were contrasted using the C_2_ were: populations of the invasive American road and the native ones (C_2_^Am^), populations of the invasive European road and the native ones (C_2_^Eu^), invasive populations and the native ones (C_2_^WW^). This choice was made according to the invasion roads inferred using microsatellite markers (Fraimout et al. 2017), the populations structure assessed with SNP/InDel markers called in these samples (Olazcuaga et al. 2020) and the population structure assessed here with TE markers (supplementary fig. S5). For each putatively adaptive insertion, gene vicinity in a 1 kb region centered on the insertion was investigated as described in the paragraph “Genome Wide Association Study with TE family abundance”. The presence of the insertion in a region of selective sweep was assessed using Tajima’s D. For the 22 populations, we investigated if the Tajima’s D estimated in the 1 kb window containing this insertion was inferior to the quantile 0.05 of Tajima’s D distribution in this population. More precisely, to prevent for a difference between autosome and X chromosome, autosomal insertions were compared to the autosomal Tajima’s D distribution and X-linked insertions to the X chromosome Tajima’s D distribution (with autosomal and X-linked contigs as defined in Paris et al. (2020)). We also checked if the insertion was close to SNPs/InDels previously identified as potentially adaptive during *D. suzukii* invasion (considering a maximum distance of 5 kb) (Olazcuaga et al. 2020).

## Acknowledgements

This work was supported by the French National Research Agency (ANR-16-CE02-0015-01 – SWING) and performed using the computing facilities of the CC LBBE/PRABI. We sincerely thank C. Mermet-Bouvier for technical help. We are also grateful to B. Prud’homme and F. Sabot for constructive discussion about this article.

## Supplementary data

Supplementary data are available at Molecular Biology and Evolution online.

