## Supplementary Material for "The worldwide invasion of *Drosophila suzukii* is accompanied by a large increase of transposable element load and a small number of putatively adaptive insertions"

#### Supplementary figures

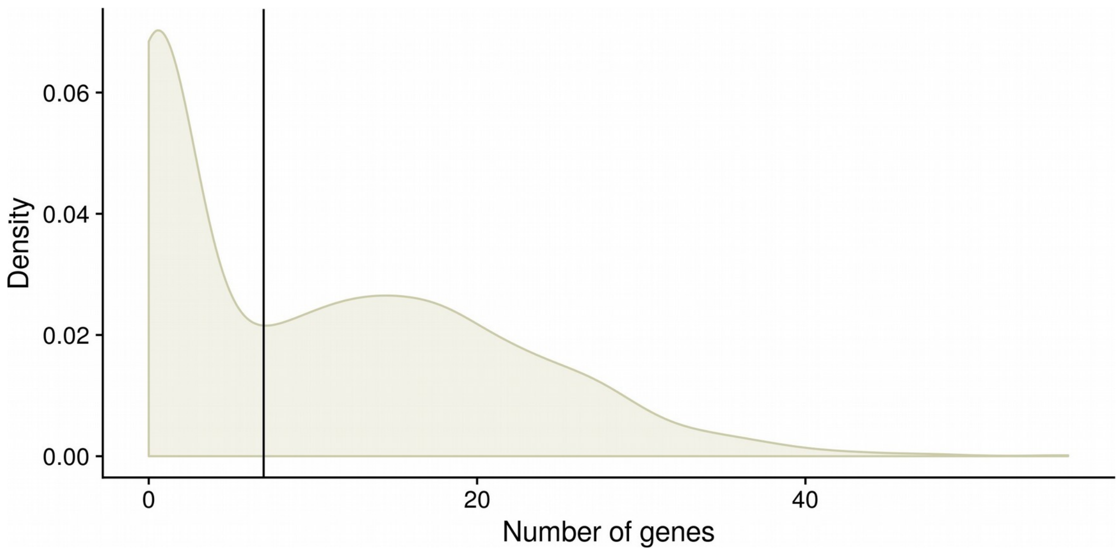


**Figure 1:**

Distribution of the number of genes per 200 kb windows in *D. suzukii* assembly. The vertical line corresponds to x=7.

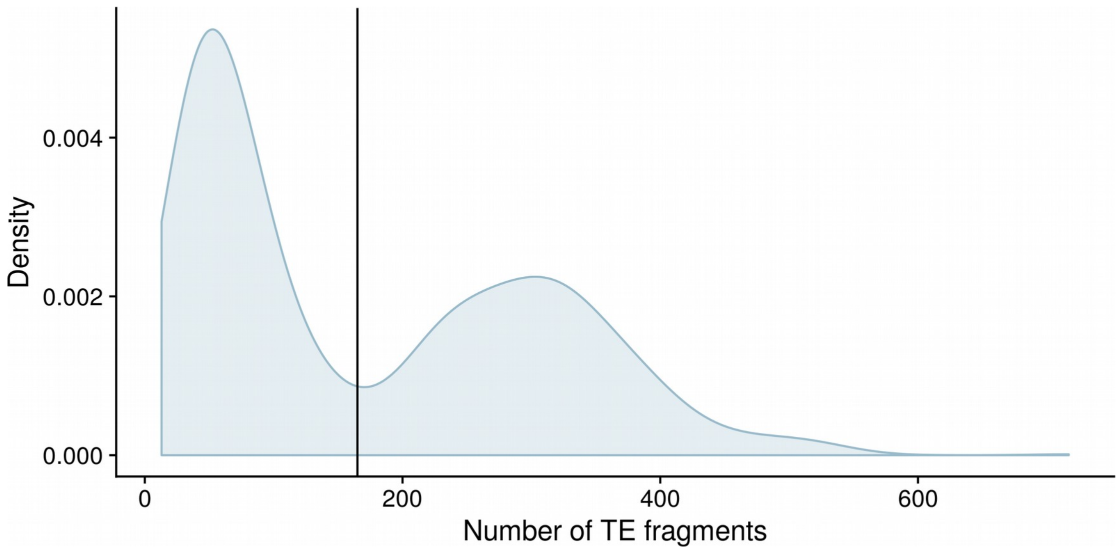


**Figure 2:**

Distribution of the number of TEs fragments per 200 kb windows in *D. suzukii* assembly. The vertical line corresponds to x=165

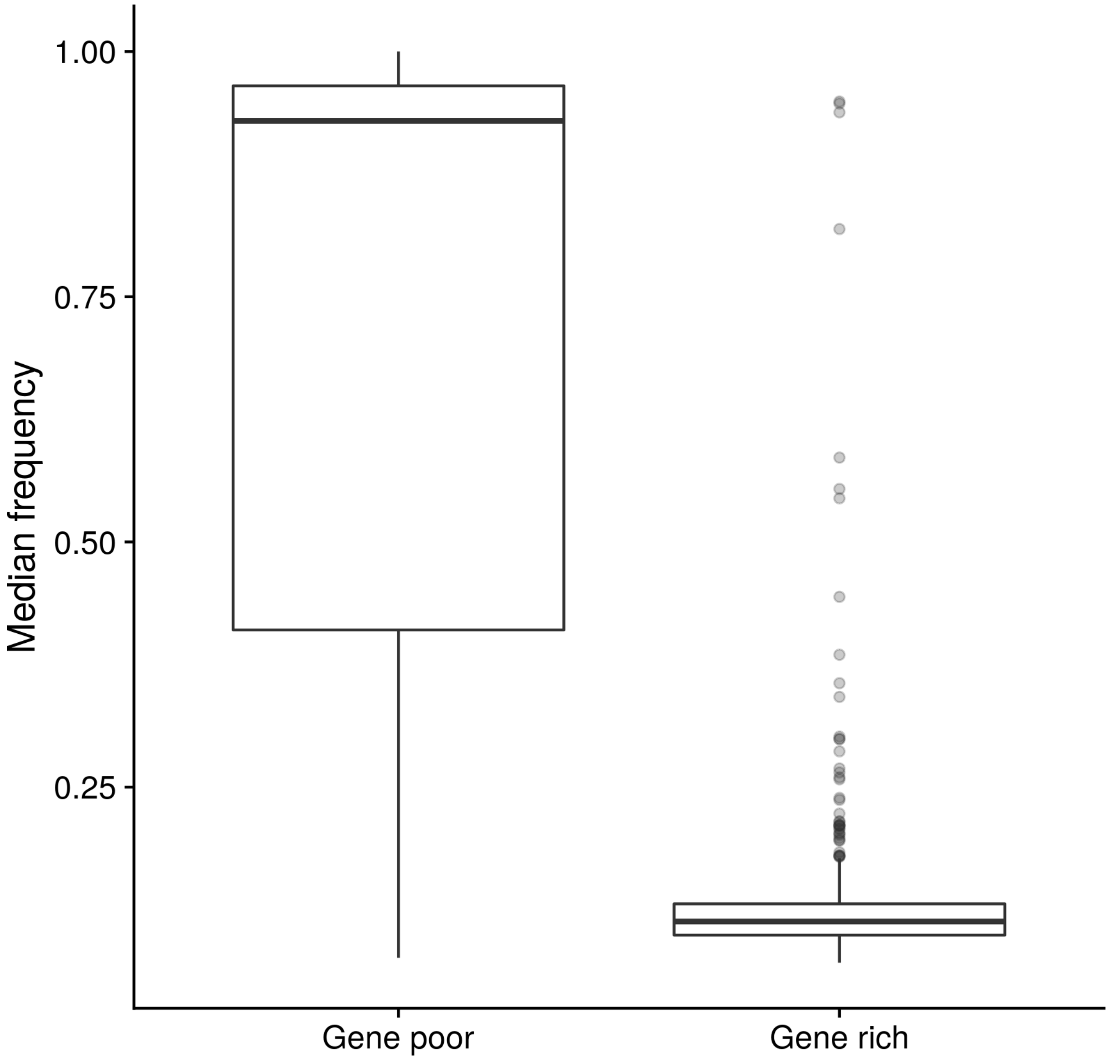


**Figure 3:**

Distribution of the median TE insertion frequency per 200 kb windows in *D. suzukii* assembly for gene-poor (< 7 genes per Mb) or gene-rich (≥ 7 genes per Mb) windows. Frequencies were estimated in Watsonville reference population.

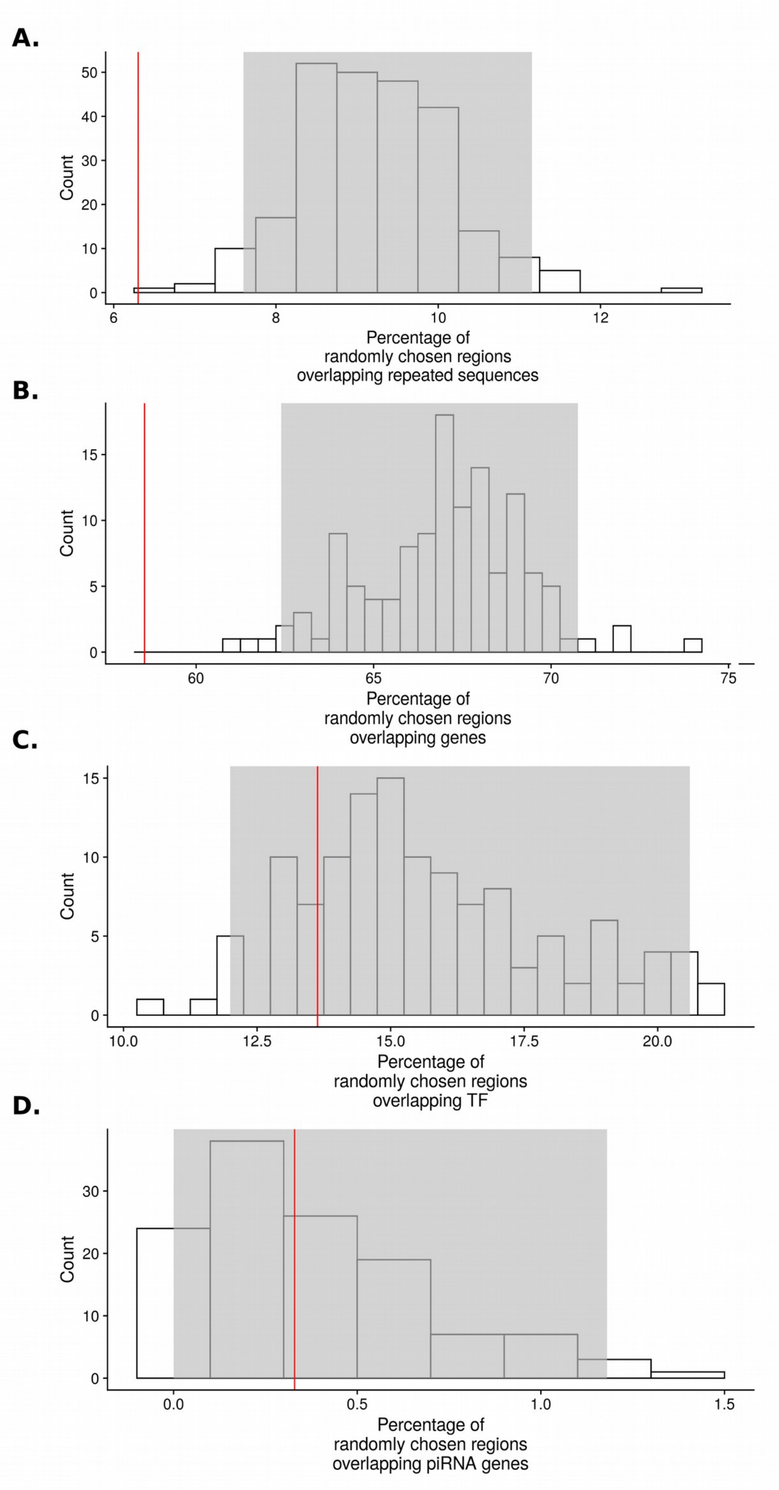


**Figure 4:**

Distribution of the percentage of randomly chosen regions surrounding SNPs in *D. suzukii* assembly and overlapping: A. repeated sequences; B. genes; C. genes of the piRNA pathway; D. genes encoding transcription factors (TFs). 250 samples of 1000 regions were used to draw the distribution A., 150 samples of 500 regions for distributions B,C and D. The gray rectangle in the background delimites the portion of the distribution between quantile 2.5% and quantile 97.5%. The vertical red lines correspond to the observed percentage for regions associated with TE abundance.

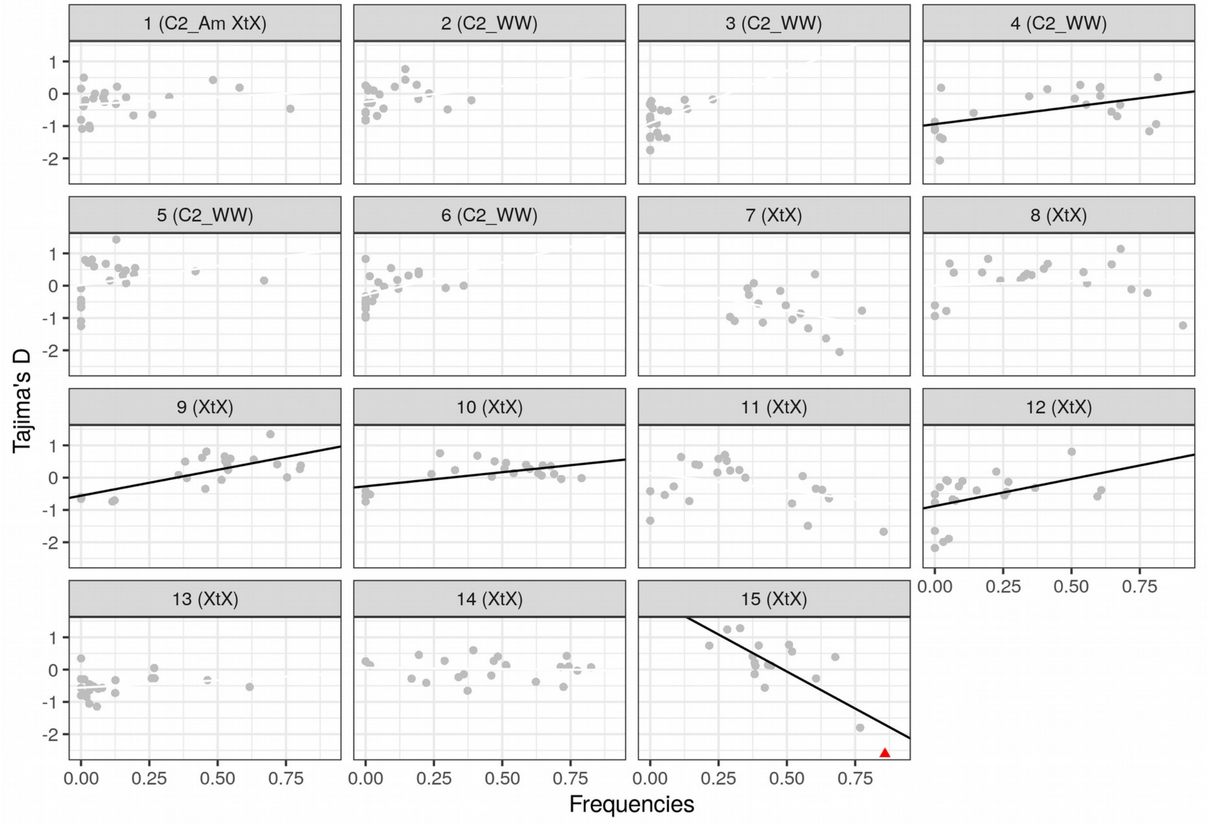


**Figure 5:**

Correlation between insertion frequencies and local Tajima's D estimates in the 22 *D. suzukii* populations for each of the 15 putatively adaptive insertions. Each panel corresponds to one insertion and Tajima's D are estimated from the 1 kb window containing the insertion. Regression lines are drawn when linear correlations are significant (Pearson's product-moment correlation, $p<0.05$). The red dot indicates that local Tajima's D is inferior to quantile 5% in that population.

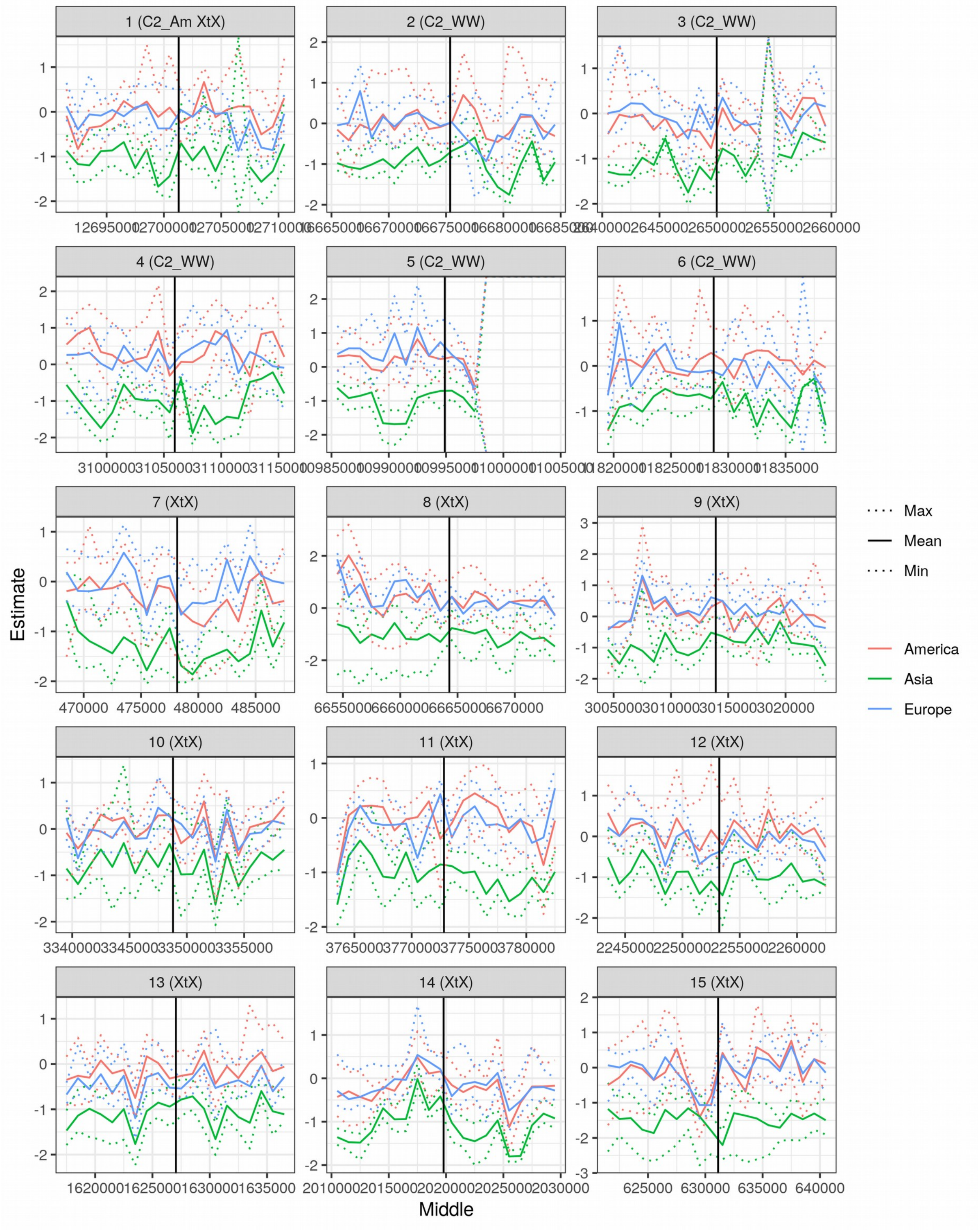
1 (C2_Am XtX)
2 (C2_WW)
3 (C2_WW)
4 (C2_WW)
5 (C2_WW)
6 (C2_WW)
7 (XtX)
8 (XtX)
9 (XtX)
10 (XtX)
11 (XtX)
12 (XtX)
13 (XtX)
14 (XtX)
15 (XtX)
Estimate
Middle
Max
Mean
Min
America
Asia
Europe

**Figure 6:**

Tajima’s D around the 15 putatively adaptative insertions. Positions along the contigs (bp) are on the x axis and TE insertions are located at the vertical black lines. Each statistics is estimated using SNPs/InDels in a 1-kb genomic window. Asian populations are in green, American in red and European in blue.

**A. Autosomes**

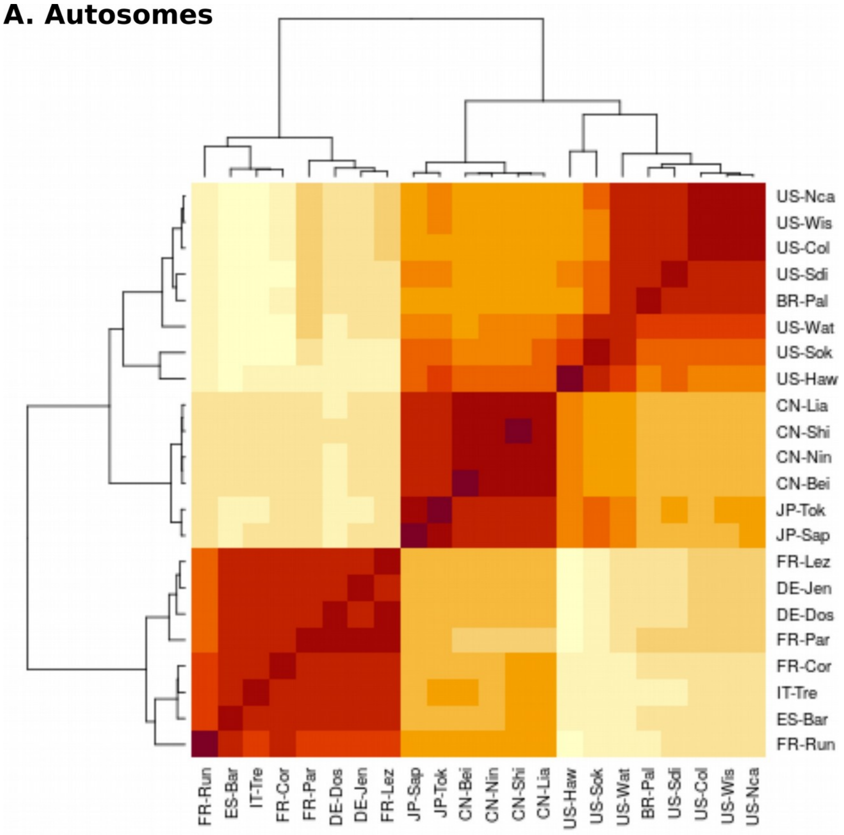


**B. Gonosomes**

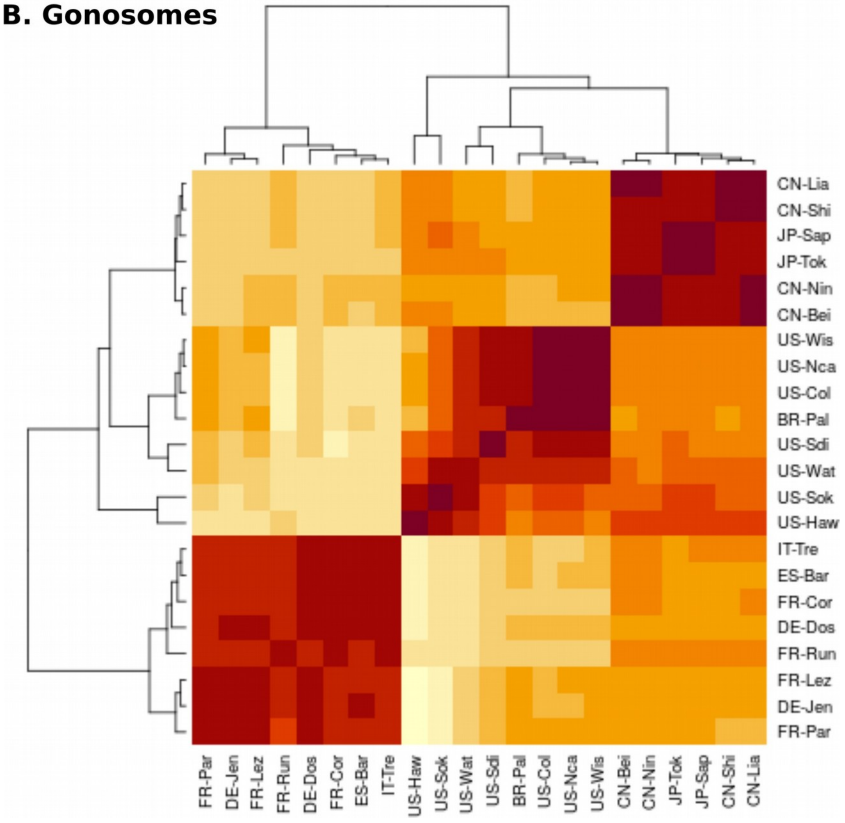

**Figure 7:**

Correlation plots of the scaled covariance matrices of population allele frequencies (Omega) among all 22 *D. suzukii* populations based on autosomal (A) and gonosomal (B) TE insertions.

### Supplementary tables

**Table 1:**

Percentage of *D. suzukii* assembly occupied by each TE superfamily

| | |
|---|---|
| CMC | 0.40 |
| Copia | 0.23 |
| CR1 | 2.41 |
| DNA | 0.05 |
| Gypsy | 13.65 |
| hAT | 0.19 |
| hAT? | 0.01 |
| Helitron | 6.95 |
| I | 2.48 |
| Kolobok | 0.05 |
| L2 | 0.48 |
| Maverick | 4.92 |
| Merlin | 0.03 |
| MULE | 0.01 |
| P | 0.03 |
| Pao | 6.44 |
| Penelope | 0.00 |
| PIF | 0.40 |
| PiggyBac | 0.03 |
| R1 | 3.24 |
| R2 | 0.03 |
| RTE | 0.13 |
| Sola | 0.00 |
| TcMar | 0.87 |
| Unknown | 4.07 |
| Zator | 0.00 |

**Table 2:**

Number of Mb of *D. suzukii* assembly attributed to each of the *D. melanogaster* chromosomes

| D. melanogaster chromosome | Mb of D. suzukii assembly |
|---|---|
| 2L | 51.9 |
| 2R | 58.8 |
| 3L | 45.6 |
| 3R | 50.0 |
| 4 | 2.6 |
| X | 31.7 |

**Table 3:**

Number of families with median of High (f ≥0.75), Intermediate (0.25≤f<0.75), or Low (f<0.25) frequency in *D. suzukii* reference population for each TE order. Only families with more than 10 insertions are considered.

| | DNA | LINE | LTR | RC | Unknown |
|---|---|---|---|---|---|
| High f. | 6 | 9 | 13 | 1 | 6 |
| Intermediate f. | 1 | 2 | 0 | 0 | 1 |
| Low f. | 18 | 21 | 19 | 5 | 17 |

#### Supplementary methods

The accuracy of the TE calling procedure was validated by a simulation work using simulaTE (Kofler 2018). As a starting point, an artificial genome devoid of TEs was created by removing masked nucleotides in a randomly selected 1 Mb chunk of the masked assembly. The resulting genome was 607,100 bp long. A population of 1000 diploid individuals each displaying 500 insertions, of frequencies ranging from 0.01 to 0.99, was generated by inserting TE sequences in the artificial genome. A reference genome, containing 250 of these insertions was also created. This artificial population was used to simulate read data corresponding to each of the 22 PoolSeq samples. For each sample, x haploid genomes were drawn according to the exact number of individuals in the sample. Reads were simulated using simulaTE and mimicking the coverage and insert size of the original sample. We used the coverage estimated in Olazcuaga et al. (2020) and the inner distance, i.e. insert size -2*Read length extracted from the ppileup file header. The standard deviation on the inner distance was set to 100 bp. The TE frequency and TE abundance pipelines described in the Materials and Methods section were then run on this dataset

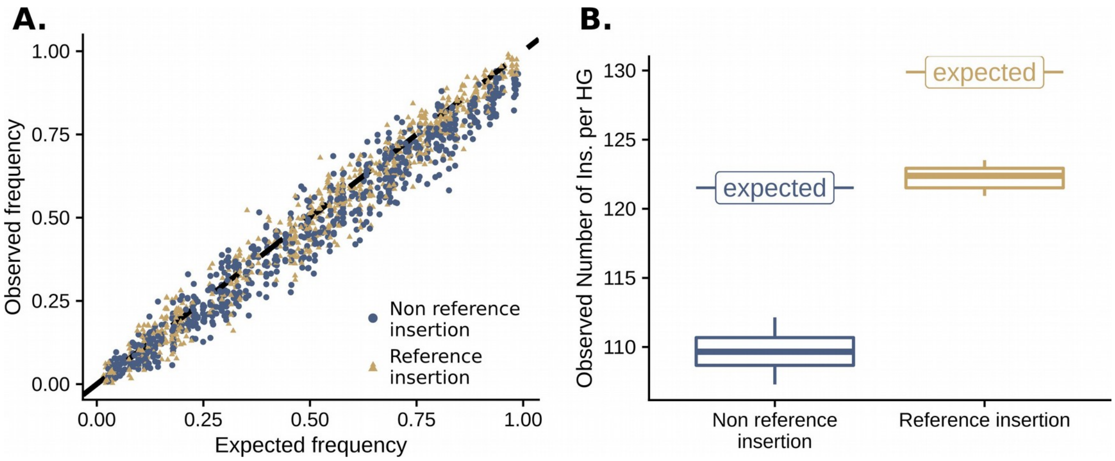


**Figure 8:**

**Validation of the TE frequency and TE abundance pipelines: expectations vs observations in a 22 samples simulated dataset mimicking the original dataset A.**

**Estimation of TE insertion frequencies**. Observed frequencies in the simulated dataset are compared to expected frequencies. Insertions absent from the reference genome are shown in blue whereas insertions present in the reference genome are in gold. **B. Estimation of TE abundance**. Distribution of the numbers of haploid insertions for the 22 simulated samples for both non reference insertions and reference insertions. The expected numbers of insertions per haploid genome (HG), 121.5 for non reference insertions and 129.9 for reference insertions, are indicated by a horizontal segment.

A run of our pipelines on a simulated dataset mimicking the original *D. suzukii* dataset indicated that our methods estimate accurately TE insertion frequencies and TE abundances (as the numbers of TE insertions per haploid genome (HG) per population) with little variation between population samples. Regarding the TE frequency pipeline, overall 10,419 TE insertions were called in the simulated dataset (Figure 8A). 10,353 of these were true positive (99.37%), 66 were false positive (0.63%). 647 insertions out of the 11,000 simulated were not recovered by PoPoolationTE2, corresponding to a false negative rate of 5.88%. The mean number of TE insertions called per sample was 473.59 (sd = 1.30), with an average of 470.59 true positives (sd = 1.30) and 3 false positives (sd = 0). The average number of false negatives was 29.40 (sd = 1.30). We found an effect of the presence of the considered insertion in the reference genome on the ability to be detected ($\chi^2$ = 32.19, df = 1, p-value = 1.4 x $10^{-8}$), insertions present in the reference genome being missed more often. The differences between expected and observed TE frequencies were poorly explained by variations in number of individuals, coverage or inner distance between samples, or their interactions ($R^2$ = 0.56%, square-root transformed Y variable). Concerning the TE abundance pipeline, the mean number of insertions per haploid genome (HG) per sample was 234.48 (sd = 1.29) for an expectation of 251.41. On average 2.59 insertions per HG (sd = 0.28) were due to false positives. A mean of 109.63 non reference insertions per HG were recovered (sd = 1.37) over the 121.52 expected. On average, 122.26 reference insertions per HG were recovered (sd = 0.84) over the 129.89 expected (Figure 8B). The difference between the mean number of insertions per HG and the expectation was higher for reference insertions compared to non-reference insertions (t = -21.924, df = 35.348, p-value < 2.2e-16). The difference between the observed mean number of insertions per HG and the expectation was poorly explained by differences in number of individuals, or coverage or inner distance between samples, or their interactions (F-statistic=1.509, df=7-14, p-value=0.24, $R^2$ = 0.43%, adjusted $R^2$=0.145, ).


1. Kofler R. SimulaTE: Simulating complex landscapes of transposable elements of populations. Bioinformatics. 2018;34: 1439. doi:10.1093/bioinformatics/btx832

2. Olazcuaga L, Loiseau A, Parrinello H, Paris M, Fraimout A, Guedot C, et al. A whole-genome scan for association with invasion success in the fruit fly Drosophila suzukii using contrasts of allele frequencies corrected for population structure. Molecular Biology and Evolution. [cited 12 May 2020]. doi:10.1093/molbev/msaa098
